# Tubulin Lattice in Cilia is in a Stressed Form Regulated by Microtubule Inner Proteins

**DOI:** 10.1101/596478

**Authors:** Muneyoshi Ichikawa, Ahmad Abdelzaher Khalifa, Kaustuv Basu, Daniel Dai, Mohammad Amin Faghfor Maghrebi, Javier Vargas, Khanh-Huy Bui

## Abstract

Cilia, the hair-like protrusions that beat at high frequencies to propel a cell or move fluid around the cell, are composed of radially bundled doublet microtubules. The doublet microtubule is composed of a 13-protofilament A-tubule, a partial 10-protofilament B-tubule and microtubule inner proteins (MIPs) inside the tubulin lattice. In this study, we present the near-atomic resolution map of the *Tetrahymena* doublet microtubules. The map demonstrates that the network of microtubule inner proteins is weaving into the tubulin lattice, forming an inner sheath of proteins. In addition, we also obtain the tubulin lattice structure with missing MIPs by Sarkosyl treatment. In this structure, the tubulin lattice showed significant longitudinal compaction and lateral angle changes between protofilaments. These results are evidence that the binding of MIPs directly affects and stabilizes the tubulin lattice. It is also suggested that the doublet microtubule is an intrinsically stressed filament and this stress could be exploited in the regulation of ciliary waveforms.

## Introduction

Microtubules are tubular structures composed of protofilaments (PFs) of α- and β-tubulin heterodimers in eukaryotes and are responsible for structural support, tracks in intracellular transport and organization of organelles. In the cilia, nine doublet microtubules (doublets) are radially bundled to form an axonemal structure. The doublet is composed of a complete 13-PF A-tubule and an incomplete 10-PF B-tubule (Fig. 1A). The doublet is the scaffold where ciliary proteins, such as axonemal dyneins and radial spokes, are periodically docked (1). These proteins are important to initiate and regulate the bending motion of the cilia. The doublet also serves as the tracks for motor proteins kinesin-2 and dynein-2 carrying intraflagellar transport cargoes towards the distal tip and back to the base of the cilia (2). Defects in ciliary proteins cause abnormal motility and function, hence, leading to cilia-related diseases, such as primary cilia dyskinesia and Bardet-Biedl syndrome (3).

**Fig. 1.**
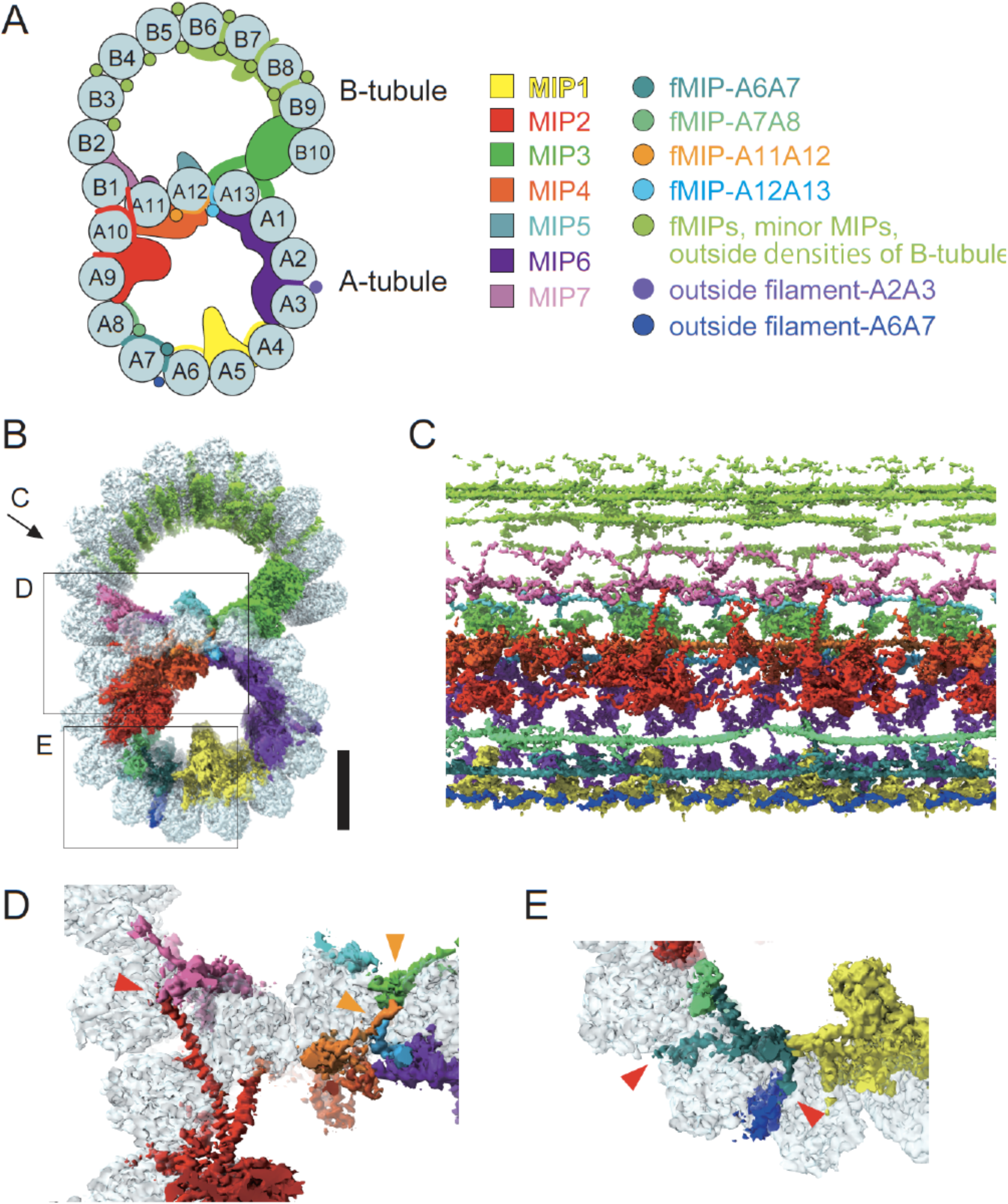
Network of the MIPs are woven into the tubulin lattice. Schematic cartoon of the doublet from *Tetrahymena* viewed from the tip of the cilia. PF numbers are shown, and MIPs are colored as the right panel. (B) Surface rendering of the 48-nm repeating unit of the doublet [colors according to (A)]. Scale bar, 10 nm. Views of (C-E) is indicated in (B). (C) The weaving network of MIPs inside the tubulin lattice. Tubulin densities were removed for clarity. (D and E) Insertions of the MIPs woven into the tubulin lattice. Mostly α-helical branches from MIP densities were going into (orange arrowheads) or through (red arrowheads) the tubulin lattice the tubulin lattice. The branch from MIP2 interacts with MIP7 densities and branches from fMIP-A6A7 also reach to the external surface or outer filament-A6A7 as shown by red arrowheads.

In contrast to the singlet microtubules (singlets) that show cycles of growth and shrinkage called dynamic instability (4), the doublets are highly stable both *in vivo* and after purification (5). In the lumen of the doublet, microtubule inner proteins (MIPs) bind periodically with 48-nm repeating unit to the tubulin lattice as shown by cryo-electron tomography (6–9). Subnanometer cryo-electron microscopy (cryo-EM) structure of the doublet isolated from *Tetrahymena* cilia revealed many new MIPs forming an inner sheath inside the doublet. This inner sheath is composed of different classes of MIPs. In particular, there exists filamentous MIPs (fMIPs) composed of long α-helices running between the inner ridges of the neighbouring PF pairs (5). It is reasonable to speculate that these MIPs can exert its effects on inherent properties of the doublet such as rigidity, damage resistance and stability similar to what microtubule-associated proteins affects the properties of singlets (10).

Recently, Rib72a/b were characterized as components of MIPs inside the A-tubule of *Tetrahymena* (11) while FAP45 and FAP52 were identified as MIPs in the B-tubule in *Chlamydomonas* (10). The Rib72a/b knockout caused reduced *Tetrahymena* swimming speed. The B-tubule of *Chlamydomonas* FAP45 and FAP52 double knockout mutant were more vulnerable to depolymerization. These studies highlight the global effects of MIPs on motility and stability of cilia. The resolution of the doublet structures presented in these studies, however, was insufficient to uncover how these proteins affect the tubulin lattice. In this study, we obtained near-atomic resolution maps of the doublet and the fractionated A-tubule from *Tetrahymena* cilia to understand the influence of MIPs to the tubulin lattice.

## Results

### A weaving network of MIPs inside the tubulin lattice

To gain insights into the molecular architecture of the doublet, we obtained a 4.3 Å resolution cryo-EM map of the 48-nm repeating unit of the purified doublet from Tetrahymena cilia (Fig. S1). This map shows the details of MIPs inside the doublet (Fig. 1 B to E; Fig. S1; and Movie S1; see also the supplementary text). The MIP densities are named based on locations as in (5). At this resolution, it becomes clear that each designated MIP can contain multiple polypeptides. Instead of simply binding on the inside surface of the doublet, MIPs consist of many branches, mainly α-helical, which weave into the tubulin lattice of the doublet (Fig. 1C). In particular, some long α-helical MIP branches poke through the A-tubule into the lumen of the B-tubule (Fig. 1 D and Fig. S1, H to M). Remarkably, we also observed MIP branches lace through the A- and B-tubule to the outside, a phenomenon that has never been observed with other microtubule associated proteins. For instance, a part of MIP2 goes through the lateral gap among PF A10, A11 and B1 to the outside of the doublet (Fig. 1D and Movie S1). The branch of fMIP-A6A7 also weaves through the tubulin lattice to be in contact with outside densities (Fig. 1E), which correspond to region of outer dynein arm. Outside the B-tubule, there are a lot of densities in the groove between PF pairs from B3 to B9 (Fig. 1, A and B; Fig. S1H; and Movie S1). These densities contact the fMIPs in the B-tubule through protrusion in the tubulin lattice.

This weaving network of MIPs is more complex in the A-tubule consisting of majorly globular MIPs and some fMIPs, while the B-tubule mainly has fMIPs. As expected from the presence of the extensive weaving network of MIPs inside, the A-tubule was more stable compared with the B-tubule. After sonication, singlet A-tubules with their B-tubules physically broken were observed (Fig. S1, A and C). Treatment of the doublet with 0.2% Sarkosyl also disintegrated the B-tubule leaving the A-tubule portion as in (12) (Fig. S1, A, B, and D). This shows the importance of the MIPs in stabilizing the tubulin lattice. For a singlet, mechanical stress from repeated cycles of bending and release was shown to induce local damage in the tubulin lattice (13). This effect will be more severe in motile cilia, which beat at high frequency for long periods of motility.

Using EM images of sonicated and Sarkosyl treated fraction, we obtained two types of A-tubule structures (sonicated and Sarkosyl A-tubules) at 4.4 and 4.9 Å resolution (Fig. 2 A and B, and Fig. S1E). While the sonicated A-tubule structure retained all the MIPs inside the tubulin lattice, the Sarkosyl A-tubule structure was missing some MIP densities (Fig. 2C). Multiple densities at MIP4 area were affected by Sarkosyl treatment and a part of MIP6 was missing (Fig. 2 D and E and Fig. S2 A and B). Other MIP densities inside the Sarkosyl A-tubule were less-well resolved, suggesting they were partially removed or became flexible (Fig. 2C).

**Fig. 2.**
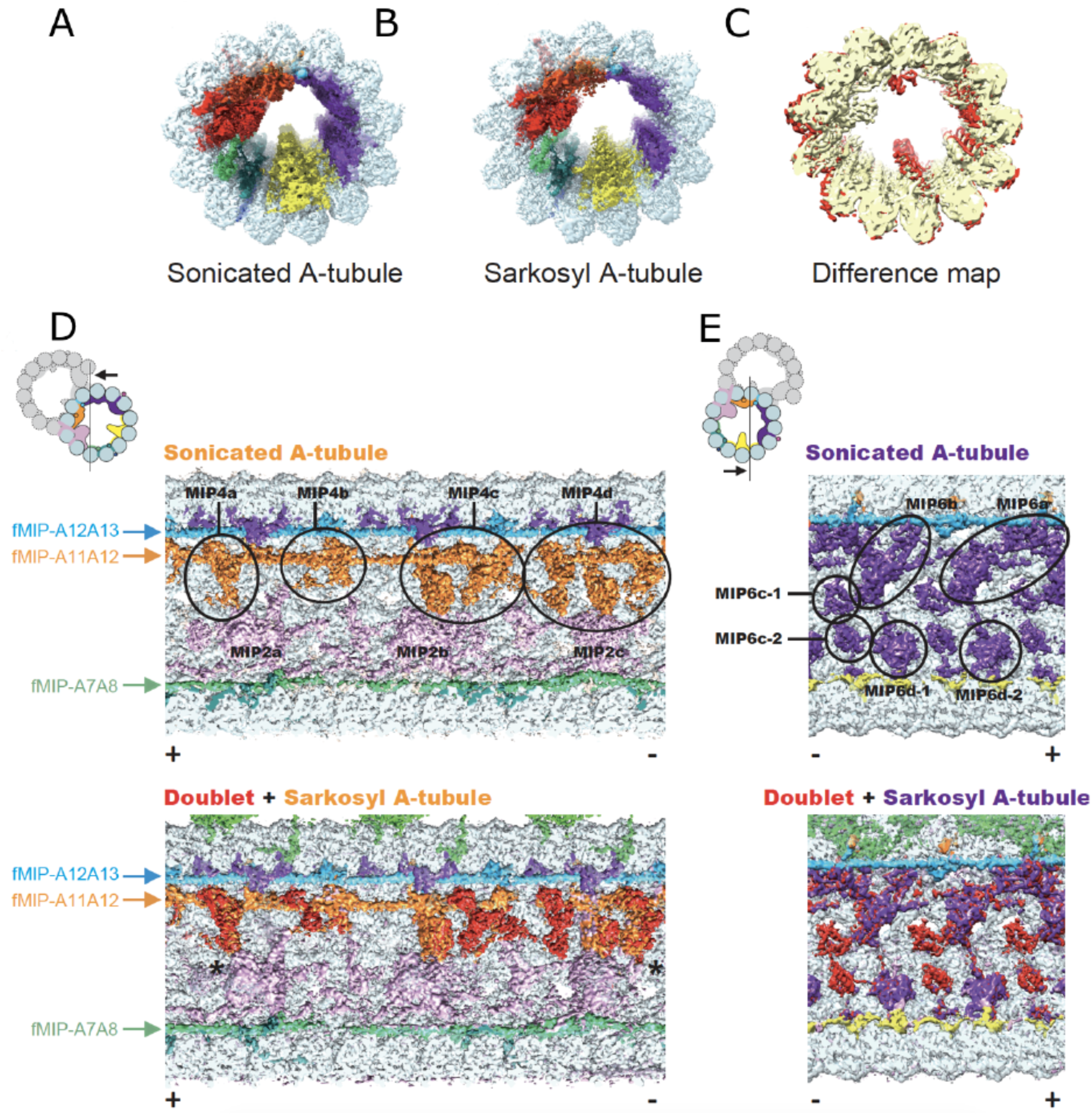
Sarkosyl treatment removes some MIPs from the doublet. (A and B) Surface rendering of the sonicated A-tubule (A) and Sarkosyl A-tubule (B) maps. (C) Difference map between the sonicated and Sarkosyl A-tubule maps. Superimposition of the two maps revealed the missing MIP densities in the Sarkosyl A-tubule map (red regions). Parts of MIP2 and MIP6 were missing in Sarkosyl A-tubule map. (C and D) Sonicated A-tubule map (top) and the overlap of doublet and Sarkosyl A-tubule maps (bottom). The MIP4 region or MIP6 region from doublet (red) are mapped onto corresponding regions from Sarkosyl A-tubule map (MIP4 in orange and MIP6 in purple). The views are indicated in the illustrations on the top left. Remaining fMIPs are indicated from the side. The coloring of MIP2 and MIP4 is different from other figures to avoid confusion (see the illustration for the coloring). Some densities at MIP4 and MIP6 regions were missing after the Sarkosyl treatment while the fMIPs appeared intact. The slight shifts in MIP4a at both + and - end sides (indicated by asterisks) are due to lateral compaction of the tubulin lattice. Polarities of microtubules are indicated by + and -.

### MIPs regulate the compaction state of the tubulin lattice

In the near-atomic resolution maps, we were able to observe the conformation of each tubulin dimer in the 48-nm unit (Fig. S1G). Recently published structures of singlets show changes in the longitudinal tubulin dimer distance depend on the nucleotide states of the β-tubulin (14–18). Stable microtubules in GTP state have elongated dimer distance while the less stable GDP-state singlet microtubules have a compacted dimer distance (19).

In the doublet, densities of GTP and GDP are observed in the α- and β-tubulin respectively (Fig. S3). Therefore, the tubulin lattice in the doublet is in GDP state. The average dimer distance of the doublet was 83.1 Å (Fig. 3 A and B, and Table S1), which is comparable to the elongated GTP- type distance in singlets (15). When MIPs are missing, the lattice of Sarkosyl A-tubule showed significant compaction and the average dimer distances became 81.1 Å (Fig. 3 B to F; Fig. S3; Table S1; and Movie S2), which is considered as a GDP-type compacted lattice (15). Since the sonicated A-tubule did not show compaction (Fig. S4I and Table S1), loss of MIPs, instead of the removal of the B-tubule caused compaction of the lattice.

**Fig. 3.**
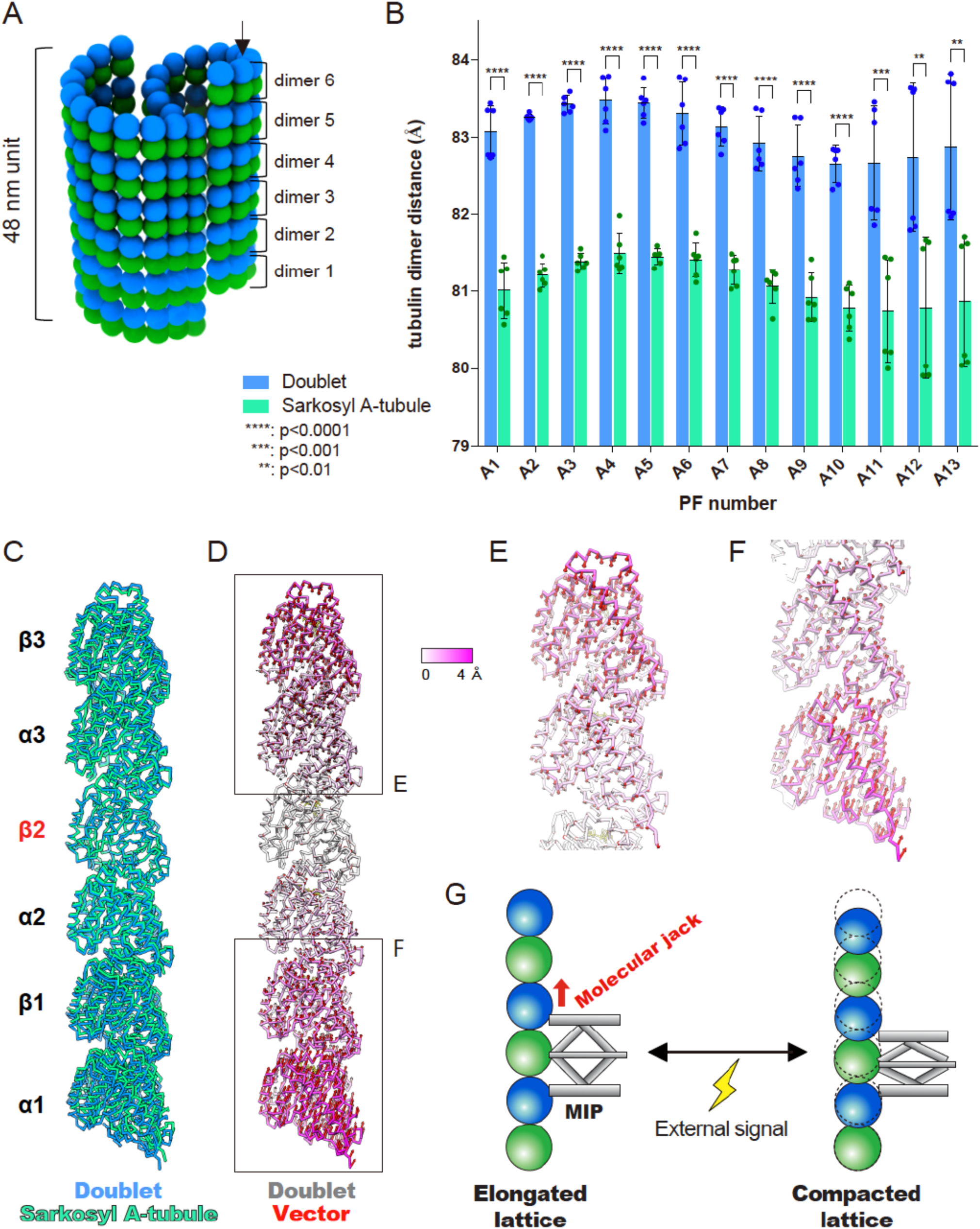
Longitudinal tubulin lattice length is regulated by MIPs. Schematics of six tubulin dimers in 48-nm repeating units of the doublet. Dimer distances were measured for each PF as in (Fig. S4A). (B) Plot of tubulin dimer distances from doublet and the Sarkosyl A-tubule. Raw measurement (n=6) and mean value with standard deviation for each PF are shown. The average value of each PF from Sarkosyl A-tubule shows lateral compression of ~2 Å. Some PFs showed a bimodal distribution of dimer distances. Statistical analysis was performed by multiple *t*-tests (see also table S2). (C) Comparison of tubulin models refined in PF-A12 from doublet (blue) and Sarkosyl A-tubule (green) showing a lateral compaction after missing some MIPs. Models were aligned by β2- tubulin. (D) Tubulin model of PF-A12 from doublet colored according to the degree of displacement. Vectors of the Cα displacement toward the Sarkosyl A-tubule model are shown in red. (E and F) Close-up views of the tubulins from the periphery with vectors. (G) Schematic diagram of MIP’s function in regulating tubulin lattice length. Some MIPs work as a molecular jack and change tubulin lattice according to the external signals.

There was also a variation of tubulin dimer distances among PFs and even among the same PF both in our doublet and A-tubule maps (Fig. 3B and Fig. S4, see also the supplementary text). This leads to an extremely heterogenous tubulin lattice compared with the singlet (Fig. S4B). Even within the same PF, tubulin dimer distances varied within the 48-nm unit. On the other hand, since we consistently observe this heterogeneity of tubulin lattice distance in our maps averaged from thousands of cryo-EM particles, the weaving network of MIPs is able to tightly and precisely regulate this heterogeneity within its repeating unit. The dimer distances in a few PFs (PFs-A1, A11-13) showed bimodal distributions with ~2 Å differences with an oscillatory pattern (Fig. 3B and Fig. S4C, D). The existence of the bimodal distribution in the Sarkosyl A-tubule suggests that remaining MIPs determines the tubulin dimer distances. Indeed, the branches from fMIPs- A11A12 and A12A13 were forming coiled-coils and poking into the tubulin lattice between PFs- A12 and A13 every 16-nm (Fig. S1L, M), which coincides with the repeating unit of tubulin dimer distance change of PFs-A12 and A13 (Fig. S4C).

The lattice length difference within and between PFs creates tension inside the filament; hence, the doublet is an inherently stressed filament. Previous study showed that doublets purified from sea urchin sperm flagella form helical structures depending on pH or calcium ion concentration (20). This means that the degree of inherent tension inside the doublet can be changed by external cues. There are a few MIP candidates with calcium binding domain, such as Rib72a (11) and FAP85 (21). Thus, MIPs can exploit the tubulin conformational change as a tool to modify the rigidity of the doublet and, thus, ciliary bending (Fig. 3G).

### MIPs affect the curvature of the doublet

In addition to changes in lattice compaction, we also observed changes in angles between PFs (Fig. 4A and Fig. S5A). Unlike 13-PF singlet which is a near perfect circle, the A-tubule of doublet showed squashed shape as in (5) with a variety of angles between PFs (Fig. 4A). Compared to the doublet, the PF angles from the Sarkosyl A-tubule showed significant changes at PF pairs, such as A1/A2 and A12/A13 (Fig. 4A and Table S3) where MIPs were missing (Fig. 2C and Fig. S2). The Sarkosyl A-tubule’s PF pair-A12/A13, where several MIP4 densities were lost, showed the largest change of angle even when compared to the sonicated A-tubule (Fig. 4A and Fig. S5B). The lateral curvature between PF pairs-A12/A13 of the doublet was equivalent to the curvature of the 22- PF singlet, which is highly energetically unfavorable considering *in vitro* reconstituted singlets generally take the 11-16 PF structure (22). With the loss of MIP4, the curvature shifted towards a more relaxed conformation comparable to the 18-PF singlet (Fig. 4 B and C). Therefore, some MIPs work as a molecular binder to keep the tubulin lattice at a higher degree of curvature while other MIPs serve as a molecular wedge that open neighboring PFs to flatten the degree of curvature (Fig. 4D).

**Fig. 4.**
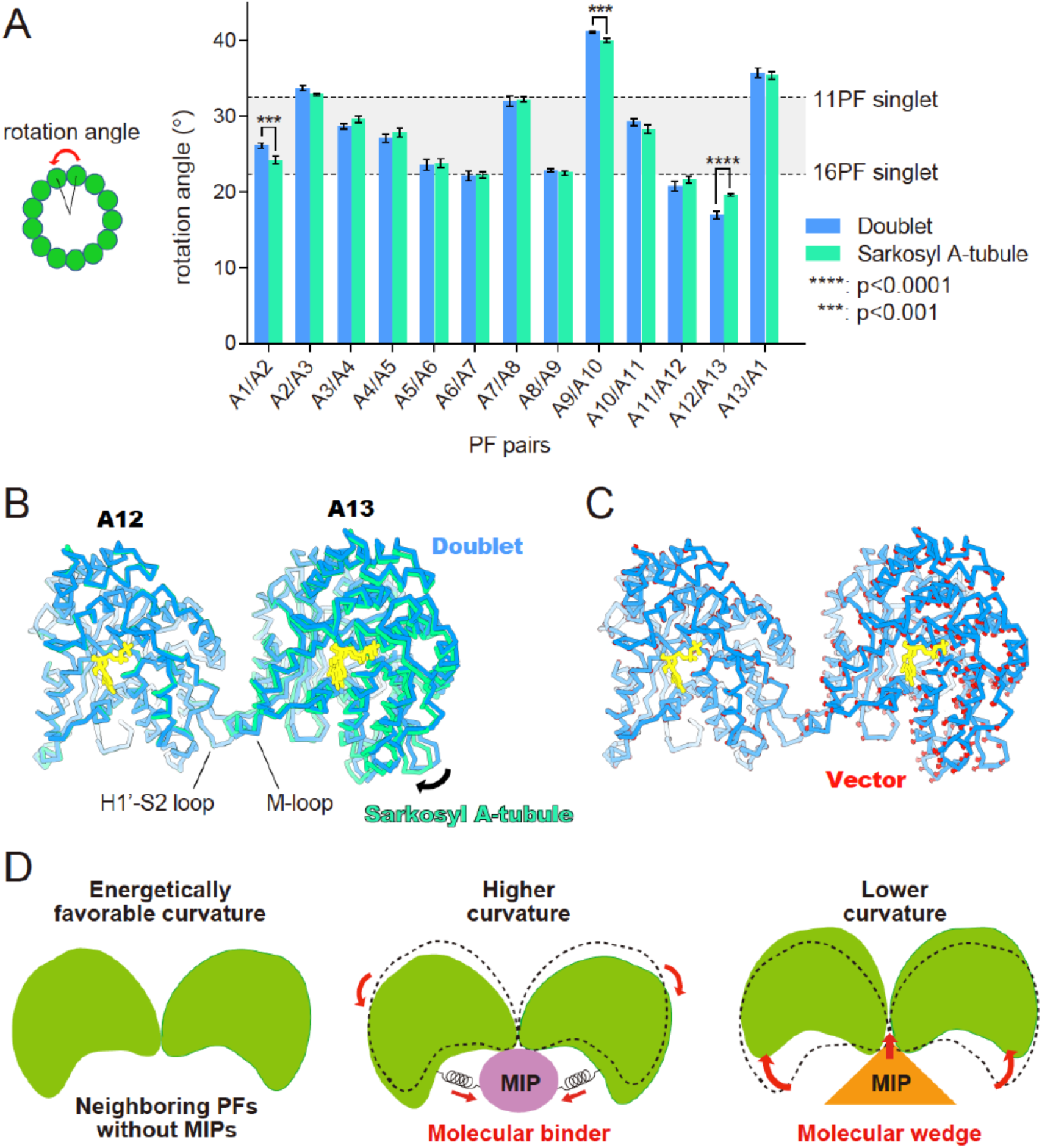
MIPs fix the tubulin lattice into extreme curvature. Plot of angles between neighboring PFs in the doublet and Sarkosyl A-tubule. Rotation angles were measured as shown in the schematic diagram on the left and mean values were plotted (see also Fig. S5A and Table S3). Error bars represent standard deviation. Multiple *t*-tests were performed to compare the mean values, and PF pairs showing *p*-values smaller than 0.001 are highlighted by asterisks (see also table S4). The gray area in the plot represents the PF pair angles commonly seen for *in vitro* reconstituted singlets (1). (B) Overlap of the models of PF pair-A12/A13 from the doublet (green) and Sarkosyl A-tubule (blue) with the tubulin unit of PF-A12 aligned reveals ~3° difference in rotation (black arrow). (C) The model of PFs-A12/A13 from the doublet with the vectors (red) of the displacement of Cα of each residue compared to the Sarkosyl A-tubule model. Nucleotides: yellow. (D) MIPs regulate the angles between PFs. Without MIPs, tubulin lattice takes energetically favorable curvature as singlets. Some MIPs work as molecular binders which holds adjacent tubulin pairs together so that it will take a higher curvature. Other MIPs work as molecular wedges and opens the PF pairs inducing a lower curvature.

The influence of MIPs on the local curvatures can be additive since the curvature of PF pair-A12/A13 after the loss of MIP4 densities is still in the energetically unfavourable range (Fig. 4A). PF pair-A9/A10 also showed a slight change in the angle without any MIPs missing (Fig. 4A). This is the location of the seam (5) (Fig. S1G) where the lateral interaction between PFs is the weakest (15). Thus, this slight angle change can be a result of the tubulin lattice accommodating the local angle changes. The global effect of local PF angle changes can be seen as a slight shift of the tubulin lattice in the map (Fig. 2 A to C).

## Discussion

Our results are the direct evidence that MIPs directly influence the tubulin lattice architecture, i.e. curvature and tubulin dimer distance from the inside. The MIPs work in a coordinated fashion to keep the doublet in a stable and distorted circle that likely facilitates the specific and proper formation of the B-tubule (5). The network of MIPs prevents the loss of tubulins and spontaneous breakage in the middle of the tubulin lattice by weaving into the tubulin lattice as an integrated layer (Fig. 5). Recently, *in vitro* reconstruction of doublet was achieved from subtilisin-treated tubulins with more flexibility of the B-tubule (23). This suggests that MIPs can limit the C-terminal conformation *in vivo* for the initial binding of the B-tubule and also stabilize the tubulin binding to the outside surface of the A-tubule. Therefore, the MIPs might play an important role in facilitating the assembly of the doublet.

**Fig. 5.**
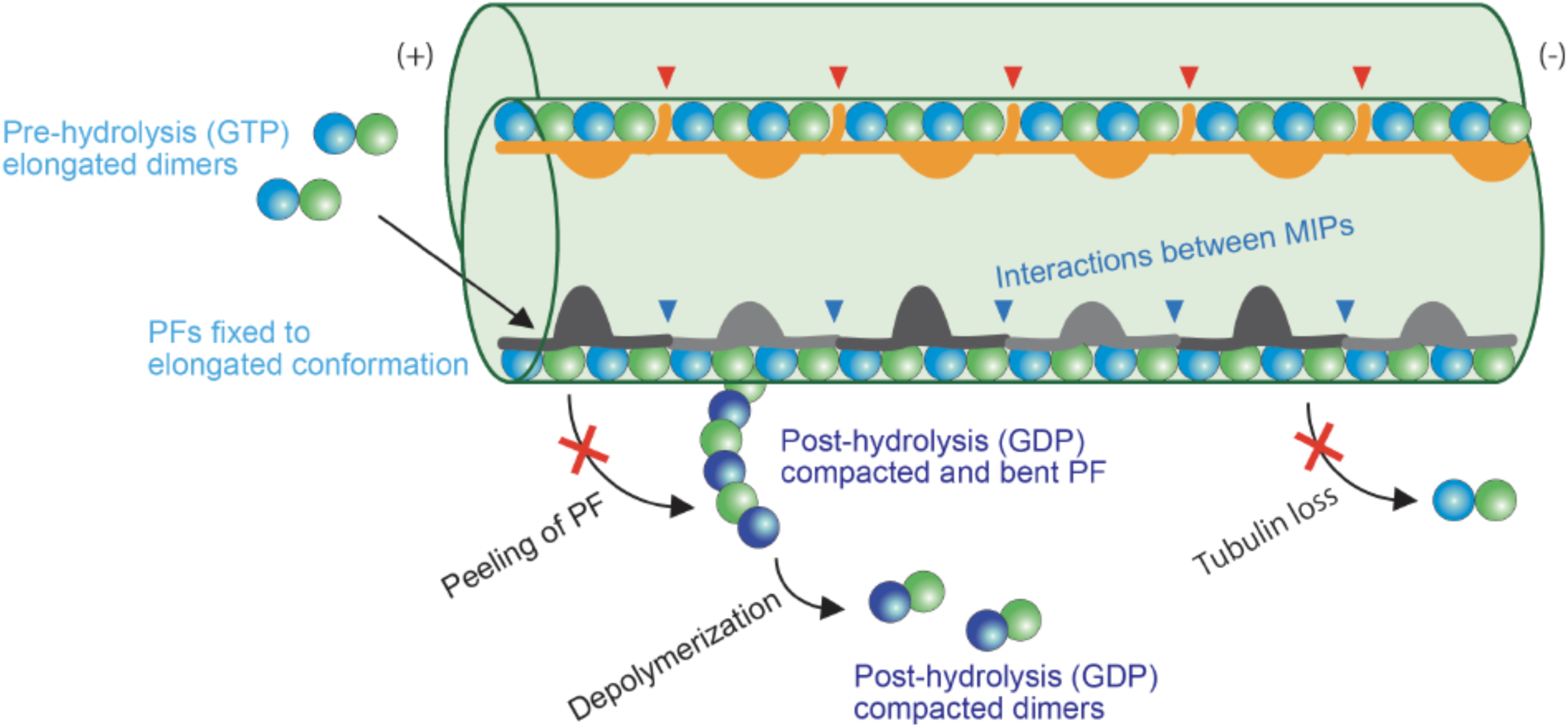
Model of stabilization mechanisms of the doublet tubulin lattice by MIPs. First, elongated tubulin dimers in GTP pre-hydrolysis state are incorporated into the tubulin lattice. This elongated and stable conformation is fixed after assembly into the doublet lattice through the interactions with the MIPs. The network of MIPs (blue arrowheads) also holds the tubulin lattice from the inside to prevent the loss of tubulin molecules or breakage at the middle part. At the plus end, MIPs prevent the peeling of PFs and depolymerization by keeping PFs in a stable and elongated conformation. Hence, double is stabilized by MIPs with several different levels to ensure that it can withstand the mechanical stress and prevent catastrophic events for the cilia. Some MIPs have insertions into tubulin lattice (red arrowheads), causing the larger inter-dimer gap and bimodal dimer distance.

Recent studies demonstrate that the elongation and compaction of the tubulin lattice of singlets plays an important role in the dynamic instability of microtubule (14–18). Binding of microtubule associated proteins can have a direct effect on the lattice compaction and, hence, microtubule dynamics (18). Our results suggest that in the doublet, the MIPs function as a molecular jack to regulate the dimer distance of the tubulin lattice in an elongated state (Fig. 3G). This points to a common mechanism where the lattice maintenance is used to regulate to microtubule stability and as the result, regulate further properties such as waveforms in the cilia.

Herein, we propose a lattice centric model for the cilia in which the doublet tubulin lattice serves as a platform to integrate the signals of binding of the MIPs and outer proteins. Binding of the MIPs leads to the local and global lattice rearrangement which affects the affinity of the outer proteins like axonemal dyneins and radial spokes. This allows the assembly of the complex axoneme in an orderly fashion (1), and thereby proper ciliary beating. The unique dimer distances among different PFs and the inside-to-outside connection can also influence the intraflagellar transport motors like kinesin-2 and dynein-2 to have different affinities and consequently use B- and A-tubule separately for transport (2).

## Acknowledgements

We thank Gary Brouhard, Susanne Bechstedt and Joaquin Ortega for helpful discussion and Kelly Sears for editing the manuscript. We thank Facility for Electron Microscopy Research of McGill University.

### Funding

This work was supported by grants from the Natural Sciences and Engineering Research Council of Canada (Discovery Grant 69462), Canada Institute of Health Research (Project Grant 388933), the Canada Institute for Advanced Research Arzieli Global Scholars Program to KHB. MI is supported by JSPS Overseas Research Fellowships. AK is a Recipient of the Al Ghurair STEM program scholarship from the Abdulla Al Gurarir Foundation for Education. We declare no competing financial interests. The electron microscopy maps are available in the EM Data Bank (www.emdatabank.org) under accession numbers EMD-XXXX to EMD-XXXX.

## Materials and Methods

### Sample preparation

*Tetrahymena* doublet microtubule fragments were prepared as in Ichikawa *et al*., (5) (fig. S1, A and B). In brief, *Tetrahymena* cells (SB255 strain) were cultured in 1L of SPP media [1% proteose peptone No.3, 0.2% glucose, 0.1% yeast extract, 0.003% ethylenediaminetetraacetic acid ferric sodium salt (Fe-EDTA)]. Cilia were isolated by dibucaine method (24) and resuspended in cilia final buffer [CFB; 50 mM 4-(2-hydroxyethyl)-1-piperazineethanesulfonic acid (HEPES), pH 7.4, 3 mM MgSO4, 0.1 mM ethylene glycol tetraacetic acid (EGTA), 0.5% Trehalose, 1 mM dithiothreitol (DTT)] containing 1 mM phenylmethylsulfonyl fluoride (PMSF). Cilia were de-membraned by adding final 1.5% NP-40, split by adding final 0.4 mM ATP, and incubated in CFB containing 0.6 M NaCl for 30 min on ice twice to remove dyneins. *Tetrahymena* doublets dialyzed against low salt buffer [5 mM HEPES, pH 7.4, 1 mM DTT, 0.5 mM ethylenediaminetetraacetic acid (EDTA)] to deplete radial spokes were fragmented by sonication and resuspended in CFB containing 0.6 M NaCl to avoid aggregation of doublet fragments.

For the Sarkosyl A-tubule, the doublets after twice 0.6 M NaCl treatment and dialysis were incubated with CFB containing 0.2% Sarkosyl to remove the B-tubule for 10 min on ice. Sonication was not performed on this sample prior to electron microscopy.

### Electron Microscopy

3.5 μl of sample of fragmented doublets (~4 mg/ml) or the Sarkosyl A-tubule (~500 μg/ml) was applied to a glow-discharged holey carbon grid (Quantifoil R2/2), blotted and plunged into liquid ethane using Vitrobot Mark IV (Thermo Fisher Scientific) at 25℃ and 100% humidity with a blot force of 3 or 4 and a blot time of 5 sec. Movies of seven frames were obtained on a Titan Krios (Thermo Fisher Scientific) equipped with Falcon II camera at 59,000 nominal magnification. The calibrated pixel size was 1.375 Å/pixel. Both datasets for the doublet and Sarkosyl A-tubule were obtained with a total dose of ~30-45 electrons/Å^2^. The defocus range was set to between −1.2 and −3.8 μm.

### Image Processing

The movies were motion corrected and dose-weighted using MotionCor2 (25) implemented in Relion3 (26) and the contrast transfer function parameters were estimated using Gctf (27). After discarding micrographs with apparent drift and ice contamination, bad contrast transfer function estimation, 7,838 micrographs for doublet and 5,179 micrographs for Sarkosyl treated A-tubule were used, respectively. The filament (doublet and A-tubule) were picked using e2helixboxer (28). Since the preparation of doublet yields both doublets and A-tubules (fig. S1A), we also picked the A-tubule from the micrographs for the doublet.

Reconstruction of the 48-nm of the doublet and A-tubule was done according to (5). In brief, 60,386 and 36,375 particles for doublet and A-tubule yielded maps of 4.7 and 4.8 Å resolution. 40,850 particles for Sarkosyl A-tubule yield 5.2 Å resolution map. After iterative per-particles-defocus refinement and Bayesian polishing in Relion 3, the resolutions of the doublet, sonicated A-tubule and Sarkosyl A-tubule improve to 4.3, 4.4 and 4.9 Å, respectively. The doublet and sonicated A-tubule maps were sharpened using Relion-3 with a B-factor of −190 and −179 Å^2^, respectively.

Since the Sarkosyl A-tubule map exhibited a slightly preferred orientation and resolution heterogeneity in the structure, we performed a local restoration and local sharpening (Javier *et al*., manuscript in preparation) to reduce artifact from preferred orientation and resolution heterogeneity. To ensure the local restoration and local sharpening does not alter the maps, we also performed the local restoration and local sharpening on the doublet and A-tubule. There was no artifact observed compared with global sharpening.

### Difference Map

To reliably identify the densities missing in the Sarkosyl A-tubule, the unsharpened maps of the sonicated and Sarkosyl A-tubule were filtered to 6 Å before performing difference mapping in Chimera. After the subtraction, the regions of difference were mapped onto the Sarkosyl A-tubule map as shown in Fig. 1H.

### Modelling

Models of the α- and β-tubulins from *Tetrahymena* (Uniprot IDs: P41351, P41352) were constructed by homology modelling in Modeller v9.19 (29) using *Sus scrofa* tubulin structure (PDB: 3JAR) as a template. The homology models of the α- and β-tubulin dimers were refined in the doublet map using Phenix (30). With our asymmetric structure of 48-nm periodicity, we were able to model, dock and refine all six individual tubulin dimers within the 48-nm structure in each individual PF of the doublet, sonicated and Sarkosyl A-tubules.

### Intra and inter-dimer distance measurement

The α- and β-tubulin can be clearly distinguished in the maps. We docked in the atomic models of the α- and β-tubulins in the map separately. The intra-dimer distance was measured as the distance between the N9 of GTP in the α-tubulin and GDP in the β-tubulin of the same tubulin dimer in Chimera. The inter-dimer distance was measured between N9 of GDP of the α-tubulin and GDP in the β-tubulin in the next tubulin dimer. The dimer distance was calculated as a sum of the intra- and inter-dimer distances.

### PF pair rotation angle (lateral curvature) measurement

The lateral curvature can be represented by the lateral rotation angle between each PF pair. The rotation angles and Z-shift between PF pairs were measured using the ‘measure’ command from UCSF Chimera (31) according to Ichikawa *et al*., (5).

### Visualization

The maps and models were segmented, coloured and visualized using Chimera (31) and ChimeraX (32).

### Data availability

The dataset analysed and raw data of the measurements are available from the corresponding author upon fair request.

## Supplementary Text

### MIPs

MIP densities have been historically named as MIP1-7 and fMIPs, it is clear that each MIP structure can contain multiple polypeptides. For example, after Sarkosyl treatment, the tip region of MIP1 was missing but the base region of MIP1 was retained (Fig. 2, A to C). MIP4 densities were widely affected by Sarkosyl treatment and part of MIP4b, c and d were lost (naming based on (5)) (Fig. 2 and S2). However, fMIP-A11A12 remained intact. This means that the globular densities and filamentous densities are composed of different polypeptides. Similarly, MIP6c region (Fig. 2) is formed by distinct polypeptide from the rest of MIP6 (MP6a, b and d) consistent with (11). In our map, MIP6c appears clearly as a two-domain structure, and thus named as MIP6c-1 and c-2 here. From the appearance of the MIP network, it is reasonable to estimate at least 30-40 polypeptides are present inside the doublet. This view is supported by the mass spectrometry of the high-salt treated axoneme of *Chlamydomonas*, showing hundreds of proteins, including proteins from the central pairs (33).

Both sonicated and Sarkosyl A-tubule maps lack B-tubule as well as MIPs binding outside the A-tubule (MIP3, MIP5 and MIP7). This suggests that most of the MIPs in the B-tubule are connected. They act as binders fixing the B-tubule onto the A-tubule and get removed together with the B-tubule.

### Outside densities

In our map, we can see the molecular ruler (34) binding to the outside furrow of PF pair-A2/A3. Even at this resolution, we did not observe any connection between the MIPs and the molecular ruler. Since the molecular ruler repeats with 96-nm periodicity (34), a 96-nm repeating unit structure might be needed to clearly describe if there is interaction or not.

In this study, we observed a few outside densities in our doublet structure even after the high-salt wash. First, there are filament-like densities in the outside wedge of PF pair-A6/A7 (shown in blue in Fig. 1 A to C; Fig. S1K; and Movie S1). These densities seem to be in contact with densities from the inside fMIP-A6A7 (red arrowheads in Fig. 1E and Fig. S1K). It is likely that the outside filament-A6A7 are a part of the outer dynein arm complex since it is located close to the binding site and these densities appear to have 24-nm periodicity, the periodicity of outer dynein arms. These inside-to-outside interactions through the tubulin lattice could coordinate the protein binding to outside and inside of doublet microtubule with different repeating units.

Outside the B-tubule, between PF pairs B3-B9, there exists outside densities (Fig. 1 A and B; Fig. S1H; and Movie S1). These densities contact the fMIPs in the B-tubule through protrusion in the tubulin lattice. Unlike the molecular ruler and outside densities of PFs A6/A7 seem filamentous, these densities do not appear filamentous. However, this is a region with lower resolution than the rest of the doublet, therefore, it is not conclusive.

### Branches of MIPs and connections to outside surface

Apart from the connections mentioned in the previous section, there are many more inside-to-outside connections. The branches of fMIP-A6A7 protrude into the tubulin lattice (Fig. 1E). These protrusions are likely to contact the outer dynein arm ruler between PF-A7/A8 binding outside *in vivo* ((35), fig. S1F).

The previously identified insertions (5) from MIP2 from inside of the A-tubule to the lumen of the B-tubule is now clearly observed as two long α-helices per 48-nm repeat (Fig. 1 B to D). These helices connect with densities from MIP7 in between PF pair B1 and A11. In addition, there is a loop-like density from MIP2 protrude outside the A-tubule and into the gap between PFs-A10 and B1 (Fig. S1I). This density is likely to be exposed to the outside of the doublet. Though MIP2 and MIP7 are connected by the branches, they are likely to be composed of different polypeptides since we see branches from MIP2 are present in A-tubule structures but MIP7 are not (Fig. 2 A and B). These outside densities or insertions could act like the mean to recruit other proteins like outside assembly scaffold, molecular motors, tubulin modifying enzymes.

### Tubulin conformation

For the compaction of the lattice, it can also be observed as shifts of remaining MIPs as well as tubulins in the maps (Fig. S2, and S4). The change in dimer distance is not accomplished through perpendicular movement to the longitudinal axis of the doublet. The tubulins are tilted in a zig-zag fashion and are compressed in an axis oblique of the longitudinal axis of the doublet (Fig. 3 E and F).

It is also noteworthy that we can measure individual tubulins in the 48-nm repeats of the doublet due to its asymmetric nature while the dimer distance measured from singlets are based on tubulin structure averaged longitudinally and laterally. In our doublet map, lattice lengths were not uniform, and every PF has a unique average dimer distance (Fig 3B; Fig. S4; and Table S1), which are extremely heterogeneous compared with the singlet (Fig. S4B). Even within the same PF, tubulin dimer distances varied within the 48-nm unit.

*Tetrahymena* tubulin lattice showed an inter-dimer distance of 41.7 ± 0.094 Å for doublet and 40.8 ± 0.159 Å for Sarkosyl A-tubule and intra-dimer distance of 41.3 ± 0.519 Å for doublet and 40.3 ± 0.486 Å for Sarkosyl A-tubule (Table S1). The deviation of intra-dimer distance was larger than that of inter-dimer distance. This agrees with other studies on microtubule structure where the inter-dimer distance is more flexible to change from nucleotide states (16). On the other hand, the compressed Sarkosyl A-tubule has shorter averaged intra-dimer distances and hence shorter dimer distances compared with singlet microtubule. The shortest dimer distance observed in the Sarkosyl A-tubule is 79.9 Å, which is shorter than any reported tubulin dimer distance (19). This could be either doublet microtubule specific feature or intrinsic feature of *Tetrahymena* tubulin. Structural analysis using doublets from other species in future studies would reveal this point.

This bimodal pattern of alternating long and shot dimer distances results in a significantly larger interaction interface at the short distance pairs compared with the long-distance pairs (Fig. S4K). In the case of Sarkosyl A-tubule, the compression of the tubulin lattice increases this interface significantly. In addition, while there is a huge variation in the dimer distance among PFs, in most cases, the differences in the dimer distance between neighbouring PFs are quite small allowing the tubulin lattice to accommodate accumulative large changes (Fig. S4B).

The lattice length of the B-tubule is slightly shorter than the A-tubule as in (Fig. S4, I and J). Previously, the doublet from sea urchin sperm was found to form spring-like structure and this was proposed to be due to shorter B-tubule lattice than A-tubule (20) consistent with our observation. The average dimer distances in the group of PFs A9-A2 and B1-B10 are 82.6 and 82.4 Å, corresponding to 0.5 and 0.9% shorter than the group of PFs A3-A8 (83.1 Å).

## Author Contributions

KHB and MI designed the experiments and MI performed cell culture, purification, vitrification of cryo grids. MI and KB performed cryo-EM data acquisition. MI, AK and KHB performed EM data processing with helps from DD and MAFM. AK performed modelling of tubulin structure. MI and KHB interpreted the data and wrote the manuscript.

## Supplementary movie legends

**Movie S1. Complex network of MIPs inside the doublet tubulin lattice.**

MIPs inside the doublet tubulin lattice were connected and forming a complex network. Some branches were reaching even outside surface.

**Movie S2. Comparison of tubulin lattice models from doublet and Sarkosyl A-tubule.** Tubulin models based on PF-A12 were morphed. After the Sarkosyl treatment, tubulin lattice showed a significant longitudinal compaction.

---

**Fig. S1.**
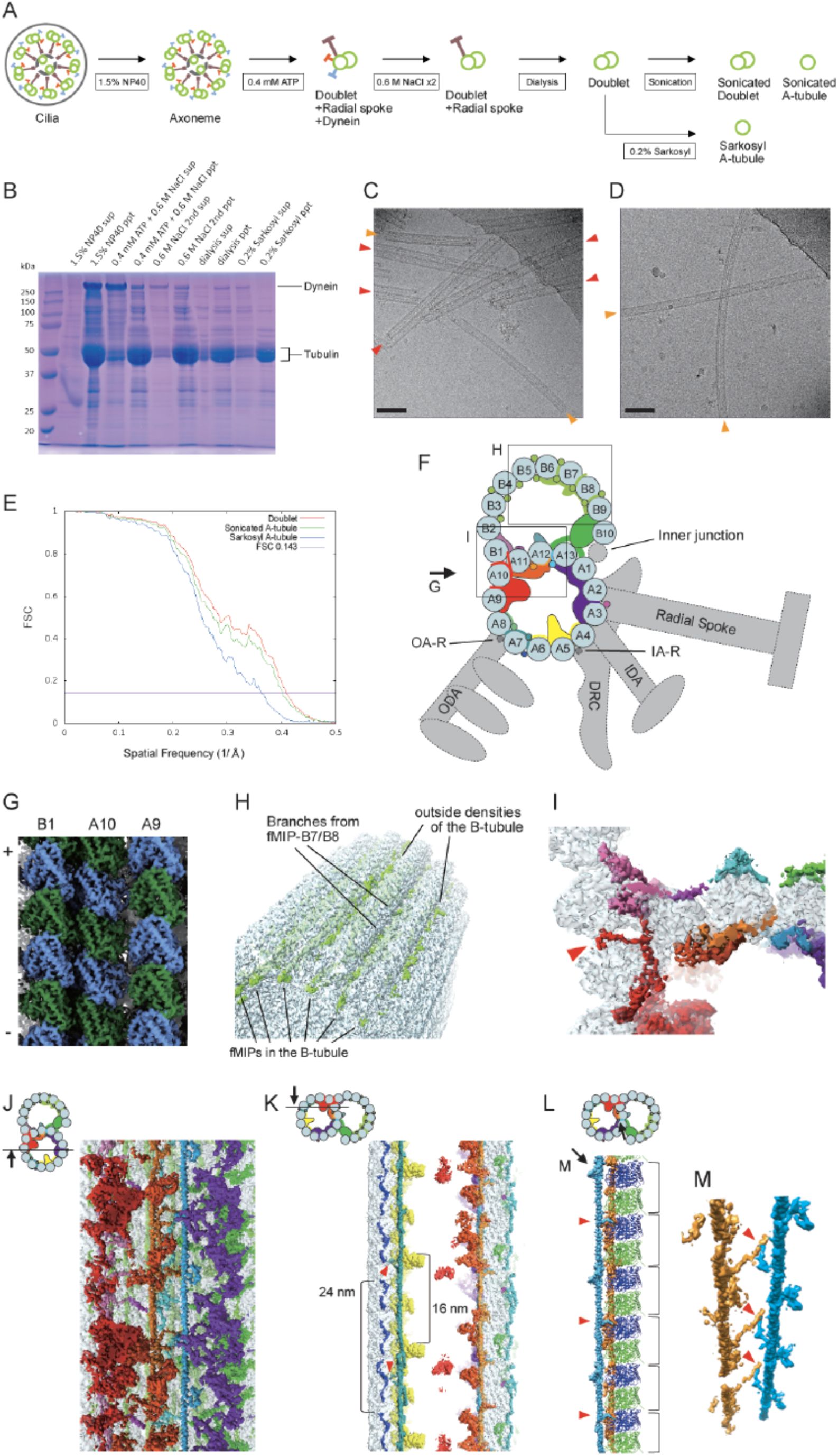
Related to doublet microtubule structure. (A) Schematics of fractionation of the axoneme in this study. Doublet microtubule were split from axoneme, and outside proteins were removed to obtain simpler sample for cryo-EM. A-tubules were obtained either sonication or Sarkosyl treatment. (B) SDS-PAGE gel of the fractionated axoneme. From the gel, Sarkosyl-treated fraction was less complex than the doublet fraction consistent with the missing densities in the EM result. (C) A typical cryo-EM image of doublet fraction showing both doublets (red arrowheads) and A-tubules (orange arrowheads) due to sonication process. (D) A representative cryo-EM image of the Sarkosyl-treated fraction showing the A-tubules (orange arrowheads). Scale bars in (C and D), 100 nm. (E) Gold-standard Fourier Shell Correlation of the doublet, sonicated A-tubule and Sarkosyl A-tubule maps. (F) Model of doublet structure with other associated structures. The parts shown by gray and dashed lines were removed in our preparation and not seen in our map. The locations of outer arm dynein-regulator (OA-R) and inner arm dynein-regulator (IA-R) are adopted from (2). (G) Surface rendering of the doublet showing PFs-A9, A10 and B1. α-tubulin is shown in green and β-tubulin in blue. (H) A view of branches from fMIPs reaching outside surface and the outside densities at B-tubule region. (I) A view of doublet map showing the branch of MIP2 (indicated by red arrowhead) going between PF pairs-A10/B1 and reaching outside of the doublet tubulin lattice. The area is the same with Fig. 1D but with a different depth. The views (G-I) are shown in (F). (J) A view of ribbon region of doublet map. fMIPs appeared as single α-helical structures running in between the inner ridges of the PF pairs-A11/A12 and A12/A13. The globular MIPs and fMIPs are connected by α-helical branches. (K) A view showing outside filament-A6A7. Red arrowheads indicate the interactions between fMIP-A6A7 and outside filament. Outside filament-A6A7 appeared as 24-nm repeating unit. (L and M) Views showing branches from fMIPs-A11A12 and A12A13. Previously, we proposed that fMIP-A12A13 has branches which pokes into the tubulin lattice every 16-nm unit (3). With higher resolution, these branches (red arrowheads) were found to be derived from both fMIPs-A11A12 and A12A13.

**Fig. S2.**
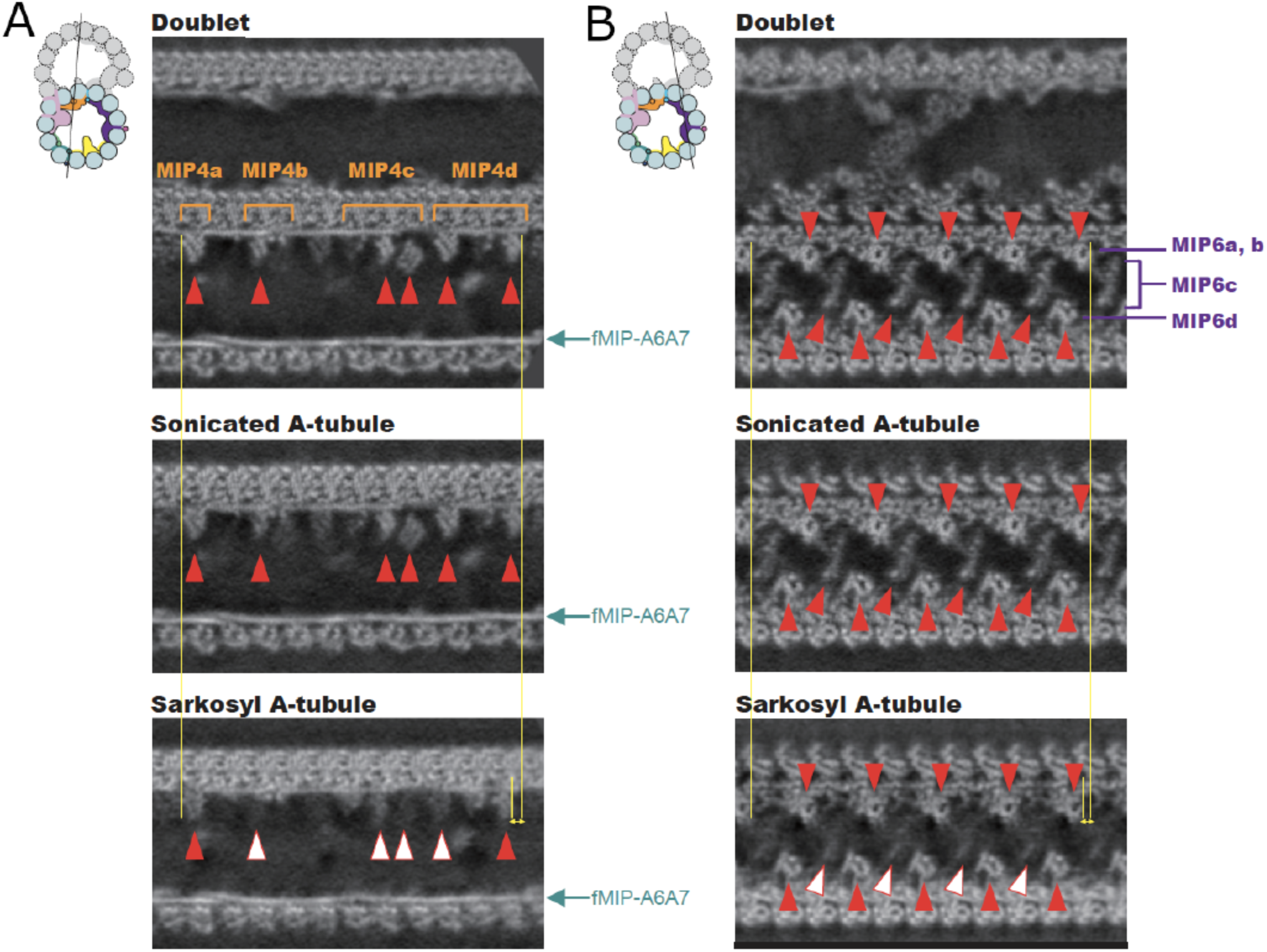
Comparison of MIPs from the doublet, sonicated A-tubule and Sarkosyl A-tubule maps. (A and B) Slices through the maps of the doublet, sonicated A-tubule and Sarkosyl A-tubule. Black lines in illustrations indicate the locations of the slices. MIP4 and MIP6 were preserved in sonicated A-tubule structure as shown by red arrowheads. Missing parts of these MIPs in Sarkosyl A-tubule map are indicated by empty arrowheads. fMIP-A6A7 densities are shown by arrows in (A). Yellow lines and double-headed arrows indicate the shifts of the MIPs in the longitudinal direction due to compaction of the tubulin lattice in Sarkosyl A-tubule.

**Fig. S3.**
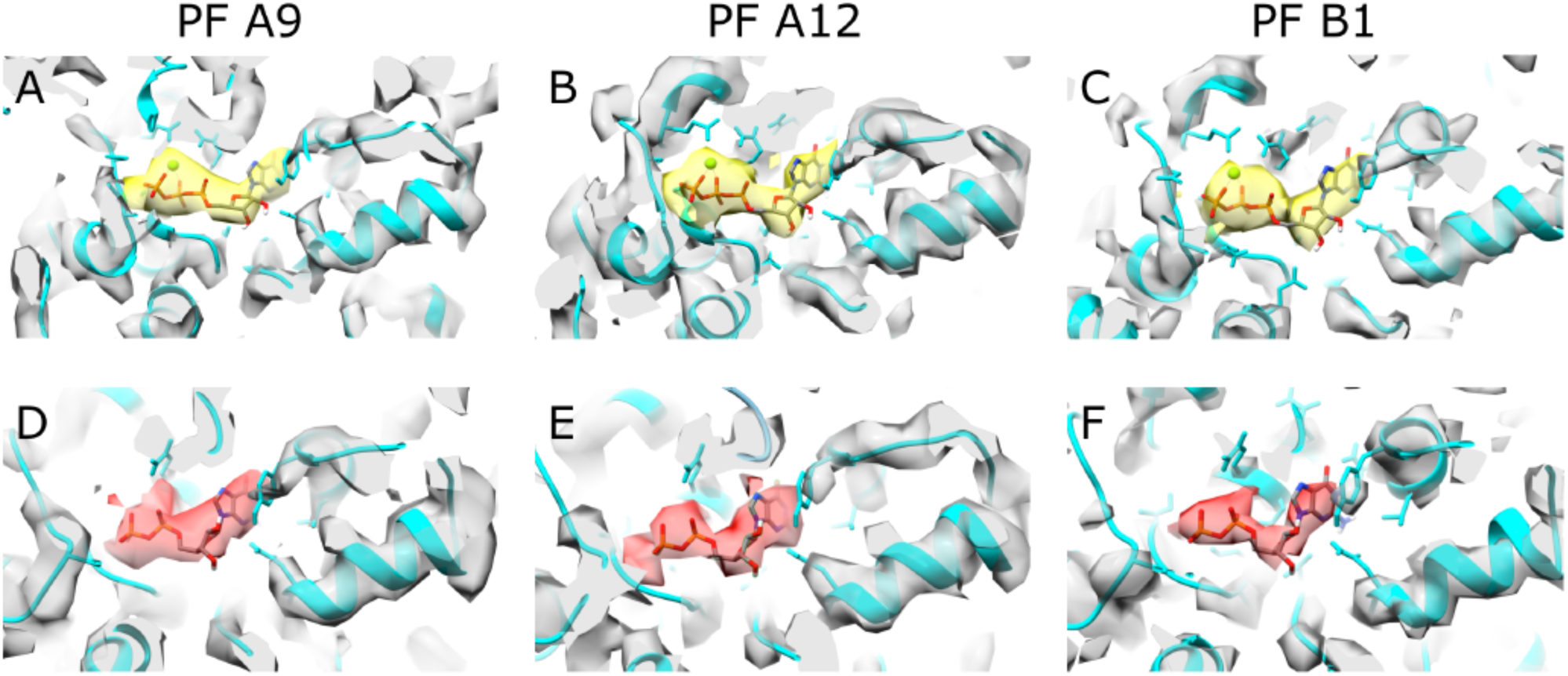
Nucleotide states in the doublet. Densities corresponding to GTP are observed in α-tubulins of PF A9 (A), A12 (B) and B1 (C) while densities corresponding to GDP are observed in β-tubulins of PF A9 (D), A12 (E) and B1 (F).

**Fig. S4.**
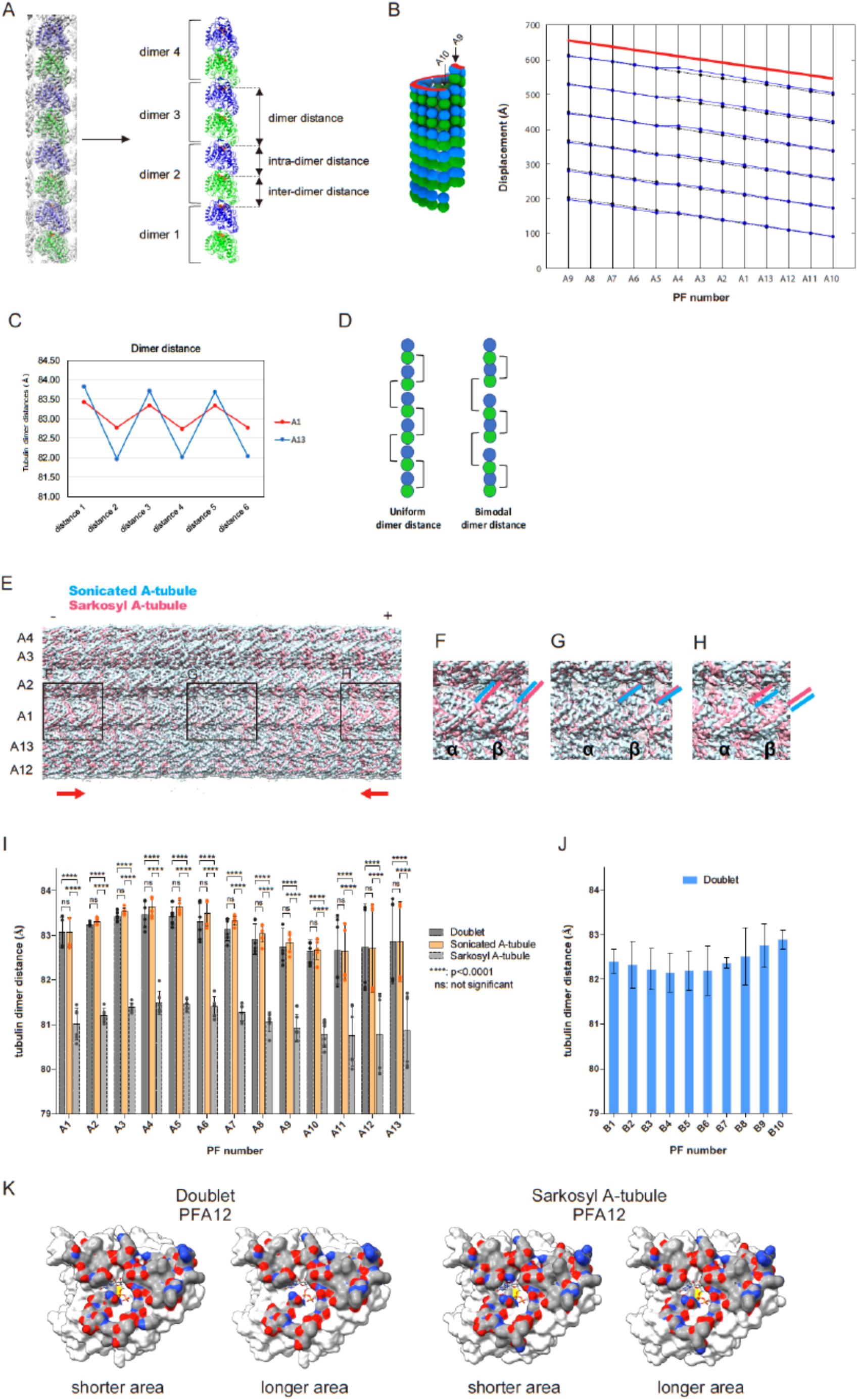
Data related to longitudinal tubulin dimer distance. (A) Details of measurement of the tubulin dimer distances. First, tubulin monomer models were fitted to each PF (left panel) and intra- and inter-dimer distances were measured as distances between nucleotide molecules (right panel). Dimer distances were obtained as sums of intra- and inter-dimer distances. Nucleotides are shown in orange. (B) Plot of the tubulin dimer distances of the A-tubule from the doublet (blue) and 13-PF singlet (1) (black). Tubulin lattice is cut and opened at the seam as in the schematic diagram. Despite having a 13-3 B-lattice as a 13-PF singlet, the A-tubule from doublet showed non-uniform tubulin dimer distances and Z-shifts. (C) Oscillation of tubulin dimer distances. The dimer distances from PFs-A1 and A13 of the doublet, where bimodal distributions were seen, were plotted in the same order as in 48-nm unit. The dimer distances oscillated with every two tubulin units (~16 nm), which coincides with repeating unit of MIPs at these regions. (D) Models of tubulin lattice with uniform dimer distance and bimodal dimer distance. In case that tubulin molecules are having same inter-dimer distances, the tubulin dimer distance will be uniform as in left panel. If tubulin molecules have alternate shorter and longer inter-dimer gaps, the tubulin dimer distance will be bimodal and oscillatory as in right panel. The distance is exaggerated for clarity and not to scale. (E-H) Comparison of the density maps of sonicated and Sarkosyl A-tubules. In the middle part, tubulin fitted well as in (G). On the other hand, as it gets closer to both ends, tubulin densities from Sarkosyl A-tubule map appeared shifted toward the middle (F and H), which means that Sarkosyl A-tubule tubulin lattice is shorter than that of sonicated A-tubule. Red arrows indicate the tubulin shift directions from both + and - ends. Locations of H12 of tubulin are indicated by pink or blue lines. (I) Plot of tubulin dimer distances from sonicated A-tubule map. Values of A-tubule from doublet map and Sarkosyl A-tubule map from Fig. 2B are shown in gray for comparison. For statistical analysis, two-way ANOVA followed by Turkey’s multiple comparison test was performed. For all PFs, changes between doublet and sonicated A-tubule were not significant (*p* > 0.01). Sonicated A-tubule also shows a bimodal distribution. (J) Plot of tubulin dimer distances of B-tubule part from the doublet. Lattice length of the B-tubule was generally shorter than the A-tubule from doublet (see also table S5). (K) Comparison of interfaces of the β-tubulin at the dimer interface. The interacting residues were colored. The interface of the shorter distance pairs at PF-A12 from doublet showed more residues involved in interactions compared with that of the longer distance pairs (1,681 vs. 1,250 A^2^). There was even more interacting interface in the shorter distance area at PF-A12 from Sarkosyl A-tubule compared with the longer distance area (1,850 vs. 1,521 Å ^2^).

**Fig. S5.**
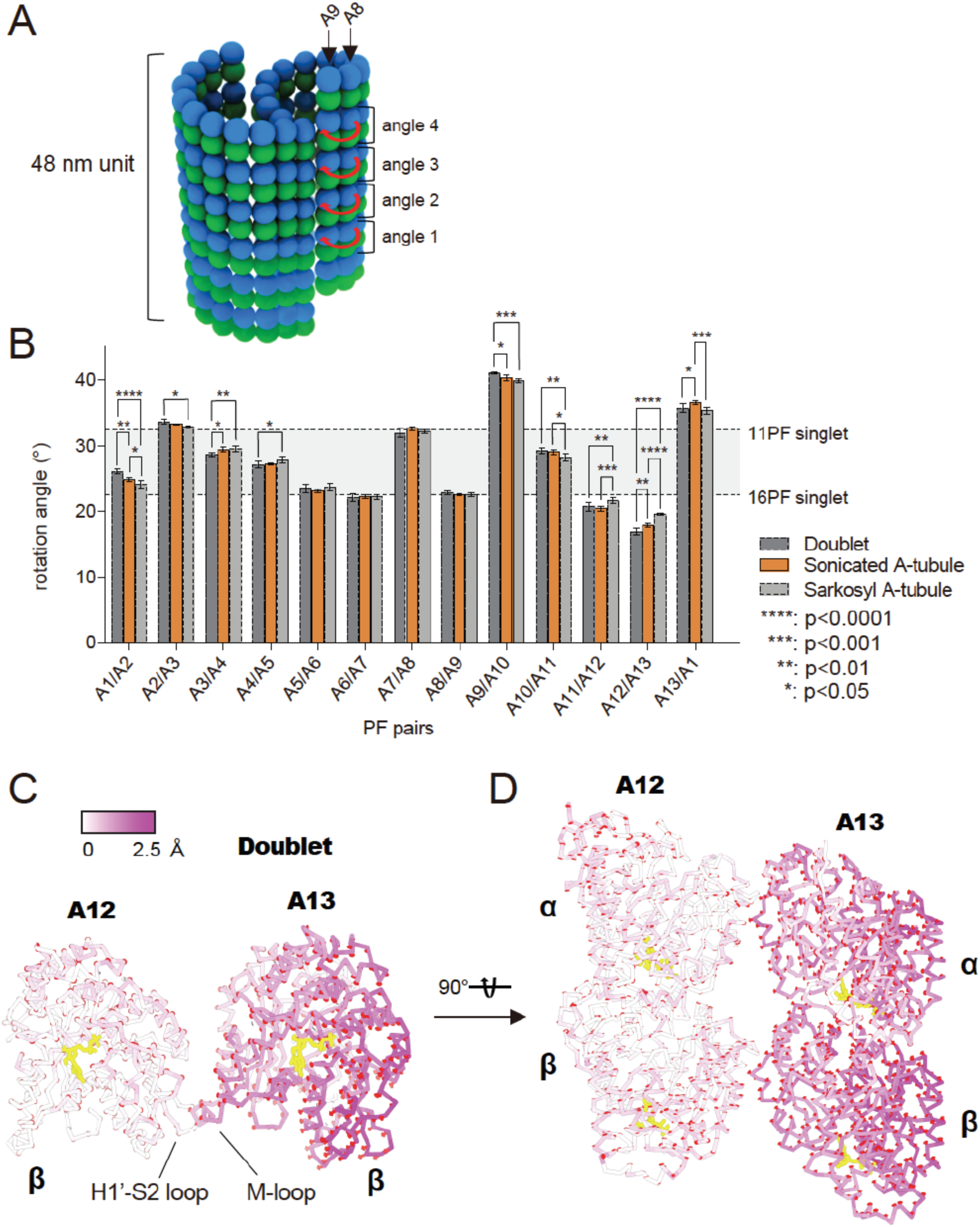
Data related to PF angle change. (A) Schematics of measurement of PF angles. Angles were measured using four tubulin pairs from each PF pair in the 48-nm unit as indicated by red arrows. PF pair-A8/A9 is shown as an example here. (B) Plots of PF angles from sonicated A-tubule map. Values of doublet and Sarkosyl A-tubule from Fig. 2A are shown in gray for comparison. Two-way ANOVA followed by Turkey’s multiple comparison test was performed for statistical analysis here. Curvatures of PFs A5-A9 where MIPs were preserved (Fig. 1, F to H) were the least affected. (C and D) Comparison of PF pair-A12/A13 models from doublet and Sarkosyl A-tubule. View in (C) is the same as Fig. 3B. The models are aligned on the tubulin dimer in PF-A12. The display model is from the PF pair-A12/A13 from the doublet and colored based on the displacement of Cα. The displacement vectors from the doublet to the Sarkosyl A-tubule are shown in red. The displacement vectors clearly show the rotation of the tubulin dimer in A13 in the Sarkosyl A-tubule. Yellow, nucleotides.

**Table S1.** Tubulin dimer distances of the A-tubule lattice.

|  | Dimer distances<br>(intra- / inter-dimer distances)<br>from doublet (Å)<br>(mean ± SD, n = 6) | Dimer distances<br>(intra- / inter-dimer distances)<br>from sonicated A-tubule (Å)<br>(mean ± SD, n = 6) | Dimer distances<br>(intra- / inter-dimer distances)<br>from Sarkosyl A-tubule (Å)<br>(mean ± SD, n = 6) |
| --- | --- | --- | --- |
| A1 | 83.1 ± 0.309<br>(41.8 ± 0.0352 / 41.3 ± 0.299) | 83.1 ± 0.313<br>(41.8 ± 0.222 / 41.3 ± 0.300) | 81.0 ± 0.333<br>(41.1 ± 0.106 / 39.9 ± 0.253) |
| A2 | 83.3 ± 0.0354<br>(41.7 ± 0.0507 / 41.5 ± 0.0636) | 83.3 ± 0.0672<br>(41.8 ± 0.0474 / 41.6 ± 0.0796) | 81.2 ± 0.143<br>(41.0 ± 0.0868 / 40.2 ± 0.0929) |
| A3 | 83.4 ± 0.0988<br>(41.8 ± 0.0242 / 41.6 ± 0.0975) | 83.5 ± 0.0696<br>(41.8 ± 0.0351 / 41.7 ± 0.0549) | 81.4 ± 0.0965<br>(40.8 ± 0.0455 / 40.6 ± 0.0965) |
| A4 | 83.5 ± 0.274<br>(41.8 ± 0.0460 / 41.6 ± 0.255) | 83.6 ± 0.195<br>(41.8 ± 0.0457 / 41.8 ± 0.191) | 81.5 ± 0.242<br>(40.8 ± 0.0511 / 40.7 ± 0.215) |
| A5 | 83.4 ± 0.184<br>(41.9 ± 0.0414 / 41.6 ± 0.199) | 83.6 ± 0.127<br>(41.9 ± 0.0344 / 41.8 ± 0.154) | 81.4 ± 0.0944<br>(40.7 ± 0.118 / 40.7 ± 0.198) |
| A6 | 83.3 ± 0.369<br>(41.8 ± 0.0730 / 41.5 ± 0.345) | 83.5 ± 0.271<br>(41.8 ± 0.0586 / 41.7 ± 0.224) | 81.4 ± 0.206<br>(41.0 ± 0.112 / 40.4 ± 0.216) |
| A7 | 83.1 ± 0.228<br>(41.7 ± 0.0762 / 41.4 ± 0.192) | 83.3 ± 0.0896<br>(41.8 ± 0.558 / 41.5 ± 0.102) | 81.3 ± 0.175<br>(41.0 ± 0.0914 / 40.3 ± 0.109) |
| A8 | 82.9 ± 0.319<br>(41.7 ± 0.0786 / 41.2 ± 0.253) | 83.0 ± 0.182<br>(41.7 ± 0.0337 / 41.3 ± 0.173) | 81.1 ± 0.204<br>(40.8 ± 0.0403 / 40.2 ± 0.207) |
| A9 | 82.8 ± 0.367<br>(41.7 ± 0.0653 / 41.0 ± 0.337) | 82.8 ± 0.198<br>(41.7 ± 0.0650 / 41.1 ± 0.239) | 80.9 ± 0.285<br>(40.9 ± 0.138 / 40.1 ± 0.295) |
| A10 | 82.7 ± 0.221<br>(41.6 ± 0.0545 / 41.0 ± 0.218) | 82.7 ± 0.202<br>(41.7 ± 0.0753 / 41.0 ± 0.226) | 80.8 ± 0.268<br>(40.7 ± 0.104 / 40.1 ± 0.261) |
| A11 | 82.7 ± 0.676<br>(41.7 ± 0.0840 / 41.0 ± 0.649) | 82.6 ± 0.575<br>(41.7 ± 0.0583 / 41.0 ± 0.593) | 80.7 ± 0.610<br>(40.7 ± 0.0681 / 40.0 ± 0.590) |
| A12 | 82.7 ± 0.875<br>(41.6 ± 0.0900 / 41.1 ± 0.960) | 82.7 ± 0.912<br>(41.7 ± 0.0462 / 41.0 ± 0.938) | 80.8 ± 0.836<br>(40.7 ± 0.0542 / 40.1 ± 0.865) |
| A13 | 82.9 ± 0.870<br>(41.7 ± 0.0443 / 41.2 ± 0.908) | 82.9 ± 0.820<br>(41.7 ± 0.0304 / 41.1 ± 0.836) | 80.9 ± 0.772<br>(40.7 ± 0.0775 / 40.1 ± 0.833) |
| All* | 83.1 ± 0.540<br>(41.7 ± 0.0937 / 41.3 ± 0.516) | 83.1 ± 0.546<br>(41.8 ± 0.0810 / 41.4 ± 0.516) | 81.1 ± 0.485<br>(40.8 ± 0.158 / 40.3 ± 0.483) |

\*For all PF results, mean values with SD calculated from all PFs (n = 78) are shown

**Table S2.** *p*-values from multiple *t*-tests of tubulin dimer distances.

|  | *p*-values<br>doublet vs<br>sonicated A-tubule | *p*-values<br>doublet vs<br>Sarkosyl A-tubule | *p*-values<br>sonicated A-tubule vs<br>Sarkosyl A-tubule |
| --- | --- | --- | --- |
| A1/A2 | 0.977 | 0.00000144 | 0.00000147 |
| A2/A3 | 0.129 | 0.0000000000289 | 0.0000000000441 |
| A3/A4 | 0.0729 | 0.0000000000155 | 0.0000000000212 |
| A4/A5 | 0.266 | 0.000000272 | 0.0000000252 |
| A5/A6 | 0.0723 | 0.00000000105 | 0.0000000000283 |
| A6/A7 | 0.380 | 0.00000151 | 0.0000000818 |
| A7/A8 | 0.110 | 0.0000000495 | 0.000000000478 |
| A8/A9 | 0.447 | 0.000000692 | 0.0000000164 |
| A9/A10 | 0.651 | 0.00000492 | 0.000000216 |
| A10/A11 | 0.857 | 0.000000284 | 0.000000183 |
| A11/A12 | 0.966 | 0.000842 | 0.000491 |
| A12/A13 | 0.966 | 0.00483 | 0.00594 |
| A13/A1 | 0.983 | 0.00319 | 0.00270 |

**Table S3.** Angles between PFs.

| | Doublet (°)<br>(mean ± SD, n = 4) | Sonicated A-tubule (°)<br>(mean ± SD, n = 4) | Sarkosyl A-tubule (°)<br>(mean ± SD, n = 4) |
| --- | --- | --- | --- |
| A1/A2 | 26.1 ± 0.341 | 24.9 ± 0.231 | 24.2 ± 0.448 |
| A2/A3 | 33.7 ± 0.303 | 33.3 ± 0.0913 | 32.9 ± 0.159 |
| A3/A4 | 28.7 ± 0.273 | 29.4 ± 0.295 | 29.6 ± 0.410 |
| A4/A5 | 27.1 ± 0.426 | 27.2 ± 0.125 | 27.8 ± 0.458 |
| A5/A6 | 23.6 ± 0.571 | 23.2 ± 0.211 | 23.8 ± 0.505 |
| A6/A7 | 22.2 ± 0.542 | 22.3 ± 0.245 | 22.2 ± 0.358 |
| A7/A8 | 32.0 ± 0.596 | 32.6 ± 0.261 | 32.2 ± 0.304 |
| A8/A9 | 22.9 ± 0.229 | 22.6 ± 0.115 | 22.5 ± 0.254 |
| A9/A10 | 41.1 ± 0.122 | 40.3 ± 0.351 | 40.0 ± 0.243 |
| A10/A11 | 29.2 ± 0.367 | 29.1 ± 0.336 | 28.3 ± 0.523 |
| A11/A12 | 20.8 ± 0.577 | 20.5 ± 0.291 | 21.7 ± 0.393 |
| A12/A13 | 16.9 ± 0.480 | 17.9 ± 0.241 | 19.6 ± 0.115 |
| A13/A1 | 35.8 ± 0.542 | 36.5 ± 0.286 | 35.4 ± 0.414 |

**Table S4.** *p*-values from multiple *t*-tests of PF-pair angles.

|  | *p*-values<br>doublet vs<br>sonicated A-tubule | *p*-values<br>doublet vs<br>Sarkosyl A-tubule | *p*-values<br>sonicated A-tubule vs<br>Sarkosyl A-tubule |
| --- | --- | --- | --- |
| A1/A2 | 0.00214 | 0.000941 | 0.0448 |
| A2/A3 | 0.0727 | 0.00820 | 0.0130 |
| A3/A4 | 0.0162 | 0.0188 | 0.647 |
| A4/A5 | 0.711 | 0.0917 | 0.0629 |
| A5/A6 | 0.314 | 0.631 | 0.102 |
| A6/A7 | 0.633 | 0.865 | 0.687 |
| A7/A8 | 0.122 | 0.569 | 0.103 |
| A8/A9 | 0.0758 | 0.145 | 0.935 |
| A9/A10 | 0.0102 | 0.000299 | 0.174 |
| A10/A11 | 0.605 | 0.0407 | 0.0669 |
| A11/A12 | 0.470 | 0.0708 | 0.00599 |
| A12/A13 | 0.0225 | 0.0000857 | 0.0000313 |
| A13/A1 | 0.0697 | 0.371 | 0.00717 |

**Table S5.** Tubulin dimer distances of the B-tubule from doublet.

| | Dimer distances<br>from doublet B-tubule (Å)<br>(mean ± SD, n = 6) |
| --- | --- |
| B1 | 82.4 ± 0.258 |
| B2 | 82.3 ± 0.476 |
| B3 | 82.2 ± 0.422 |
| B4 | 82.2 ± 0.395 |
| B5 | 82.2 ± 0.401 |
| B6 | 82.2 ± 0.502 |
| B7 | 82.4 ± 0.111 |
| B8 | 82.5 ± 0.581 |
| B9 | 82.8 ± 0.443 |
| B10 | 82.9 ± 0.195 |
| All* | 82.4 ± 0.468 |

\*For all PF results, mean values with SD calculated from all PFs (n = 60) are shown

